# Recent Bovine HPAI H5N1 Isolate Is Highly Virulent For Mice, Rapidly Causing Acute Pulmonary And Neurologic Disease

**DOI:** 10.1101/2024.08.19.608652

**Authors:** Thomas Tipih, Vignesh Mariappan, Kwe C Yinda, Kimberly Meade-White, Matthew Lewis, Atsushi Okumura, Natalie McCarthy, Chad Clancy, Emmie de Wit, Vincent J Munster, Heinz Feldmann, Kyle Rosenke

## Abstract

The highly pathogenic avian influenza (HPAI) A(H5N1) clade 2.3.4.4b viruses, responsible for the current outbreak in dairy cows in the United States, pose a significant animal and public health threat. In this study, we compared disease progression and pathology of three recent clade 2.3.4.4b isolates derived from a cow, mountain lion, and mink to a human HPAI A(H5N1) isolate from Vietnam in mice. Inoculation of C57BL/6J and BALB/c mice with all four HPAI A(H5N1) isolates resulted in comparable levels of virus replication in the lung inducing severe respiratory disease. C57BL/6J mice infected with the bovine isolate also developed high virus titers in the brain, resulting in a significant pro-inflammatory cytokine response and neurologic disease. Our findings suggest the recent bovine isolate possesses enhanced respiratory and neuroinvasive/neurovirulent properties causing fatal respiratory and neurologic disease in C57BL/6J mice.

## INTRODUCTION

Since the emergence of the highly pathogenic avian influenza (HPAI) A(H5N1) (goose/Guangdong (gs/Gd) lineage) in China in 1996 ^1^, it has evolved into 10 main phylogenetic clades ^2^. Clade 2 has been dominant and from 2005 HPAI A(H5N1) clade 2 viruses have disseminated across Asia, Europe, Africa, North and South America ^3^. Since late 2021 H5N1 clade 2.3.4.4b viruses have continuously circulated in North America causing outbreaks in wild birds and domestic poultry with increasing spillover into wild mammals ^4^. In March 2024, HPAI A(H5N1) clade 2.3.4.4b viruses were detected in dairy cows and unpasteurized milk in the United States ^5^. Thirteen human cases of HPAI A(H5N1) have so far been reported in the United States^6^. The human cases have thus far been mild, reporting mostly conjunctivitis although respiratory symptoms have been described in some cases^7,8^. All cases have been reported in persons working at dairy and poultry farms where cows or domesticated birds tested positive for HPAI A(H5N1) suggesting direct droplet cow/poultry-to-human transmission of the virus or fomite transmission through contaminated equipment. Despite evidence of mammal-to-mammal transmission, the US Centers for Disease Control currently categorizes the public health risk for human infection as low ^6^. The situation, however, could quickly change if the virus continues to acquire changes that may affect susceptibility, virulence, pathogenesis and transmission in mammalian species, specifically humans.

Prior to occurrence of HPAI A(H5N1) clade 2.3.4.4b infections in dairy cattle, outbreaks of HPAI A(H5N1) clade 2.3.4.4b viruses had been reported in mammals including farmed mink in Spain in 2022 ^9^. Disease signs in infected mink included loss of appetite, hypersalivation, depression, bleeding from the snout and neurological manifestations such as ataxia and tremors ^9^. In February 2024, a mountain lion was found dead in Montana, USA, and a HPAI A(H5N1) clade 2.3.4.4b virus was isolated from lung tissue ^10^. Similarly, natural infections in other wild terrestrial mammals have been reported ^4,11–14^ altogether highlighting irregular avian-to-mammalian spillover leading to clinical infection of varying degree in several mammal species.

The current state around HPAI (A)H5N1 creates a vulnerable condition for animal and public health in the USA and worldwide. There are many gaps to fill including countermeasure development for which animal models are critical. In this study, we compared the virulence of A/Vietnam/1203/2004 (H5N1) (VN1203), a reference HPAI A(H5N1) clade 2.3.4.4b virus, to three recent HPAI A(H5N1) isolates (A/bovine/OH/B24OSU-342/2024 (bovine), A/mountain lion/MT/1/2024 (mountain lion), and A/mink/Spain/3691-8_22VIR10586-10/2022 (mink) described above. The recent isolates are all the result of cross-species transmission likely from a wild bird source into mammalian species with the mink and bovine isolates causing significant mammal-to-mammal transmission^5,9^. We compared two potential exposure routes, nasal and oral, in C57BL/6J mice, an established model for influenza A viruses including HPAI H5N1 ^15^. We then substantiated the initial findings in a different mouse strain (BALB/c).

## RESULTS

Amino acid sequence comparison of all bovine segments with the corresponding segments of 3 additional isolates revealed the closest identity to the mountain lion (in most of the segments) followed by the mink with the most distantly related being the VN1203 isolate (**Table S1**).

### Bovine isolate rapidly induces fatal disease course

To compare virulence and pathology of the recent bovine, mink and mountain lion isolates to the reference VN1203 isolate, groups of C57BL/6J mice (n = 10) were intranasally or orally inoculated with 10^5^ TCID50 of each isolate. Four animals per group were euthanized at day 4 post inoculation (PI) for virus titer determination and histopathology. Six animals per group were monitored daily for clinical signs and survival until day 28 PI. All virus isolates induced weight loss (**Fig. 1A,B**); additional disease signs included ruffled fur and hunched posture followed by hypoactivity and rapid/labored breathing. Neurological signs, however, comprising of tremors, ataxia, hyperactivity, and circling were observed in all C57BL/6J mice inoculated with the bovine isolate. Two animals inoculated with the mountain lion isolate (1 each intranasally and orally) also exhibited minor neurologic signs of hyperactivity. Inoculation of mice with the bovine isolate by either route resulted in uniform lethality with intranasally inoculated mice succumbing by day 4 and those orally inoculated by day 6 PI (**Fig 1C,D**). Intranasal inoculation with the mink, mountain lion and reference VN1203 isolates also resulted in uniformly lethal respiratory disease. Disease progression was significantly extended reaching endpoint criteria between 4-10 days PI when compared to infection with the bovine isolate (**Fig. 1C**). Oral inoculation with the VN1203 isolate caused respiratory disease but was not uniformly lethal and similar results were found with the mink and mountain lion isolates (20-50% probability of survival) (**Fig. 1D**).

**Figure 1:**
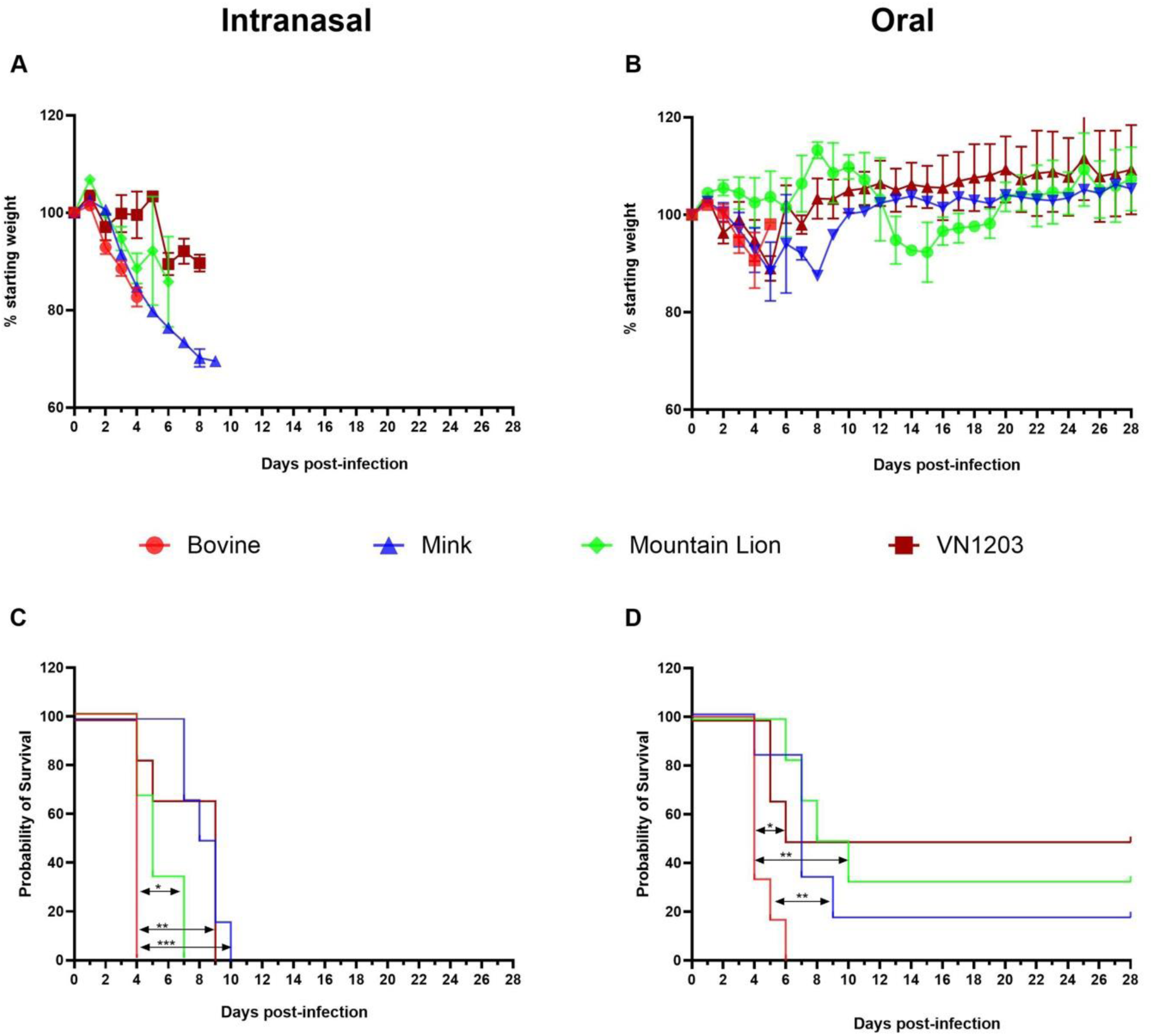
Disease progression following HPAI A(H5N1) infection. Six-week-old C57BL/6J mice (n = 10 per group) were inoculated via either intranasal or oral routes with 10^5^ TCID50 of either A/bovine/OH/B24OSU-342/2024, A/mountain lion/MT/1/2024), A/mink/Spain/3691- 8_22VIR10586-10/2022, or A/Vietnam/1203/04. Mice (n = 6) in each group were monitored for clinical signs and survival; the remaining mice (n = 4) were euthanized at a pre-determined timepoint for virology and histopathology. (A) Weight loss following intranasal inoculation (B) Weight loss following oral inoculation. (C) Survival following intranasal inoculation. Survival proportions were calculated using the Log-rank (Mantel-Cox) test. ns p > 0.05, *p < 0.05, **p < 0.01, ****p < 0.0001. (D) Survival following oral inoculation. Note, animal group sizes change over time due to animals succumbing to infection.

Survival rates were significantly lower for the bovine isolate when compared to the mink, mountain lion and VN1203 isolates (**Fig. 1C,D**). Overall, the results show that the intranasal route resulted in more severe respiratory disease than oral inoculation and that the bovine isolate was most virulent resulting in additional neurologic disease.

### Bovine isolate replicated to high titers in lung and brain tissue

To determine viral replication in organ tissues of infected C57BL/6J mice, 4 animals from each group were euthanized on day 4 PI at the onset of disease (weight loss, ruffled fur tremors, ataxia and circling). Amongst the sampled tissues, virus titers were highest in the lung samples and lowest in liver and blood samples for all the virus isolates (**Fig. 2A-H**). Consistent with neurological signs observed in C57BL/6J mice, high virus titers were found in the brains of animals infected with the bovine isolate compared to those from inoculations with the mink, mountain lion and VN1203 isolates which either were negative or low (**Fig. 2A,E**). Significantly higher viremia was detected in mice inoculated with the bovine isolate compared to those inoculated with the mink, mountain lion and VN1203 isolates of which most of the mice were negative (**Fig. 2D,H**). No significant differences in organ virus titers were observed between the oral and nasal routes. Overall, the bovine isolate demonstrated enhanced replication in organs, caused systemic infection and displayed neurotropism. In contrast, most of the mice inoculated with the mink, mountain lion, and VN1203 isolates did not show evidence of systemic infection but rather localized lung infection and respiratory disease.

**Figure 2:**
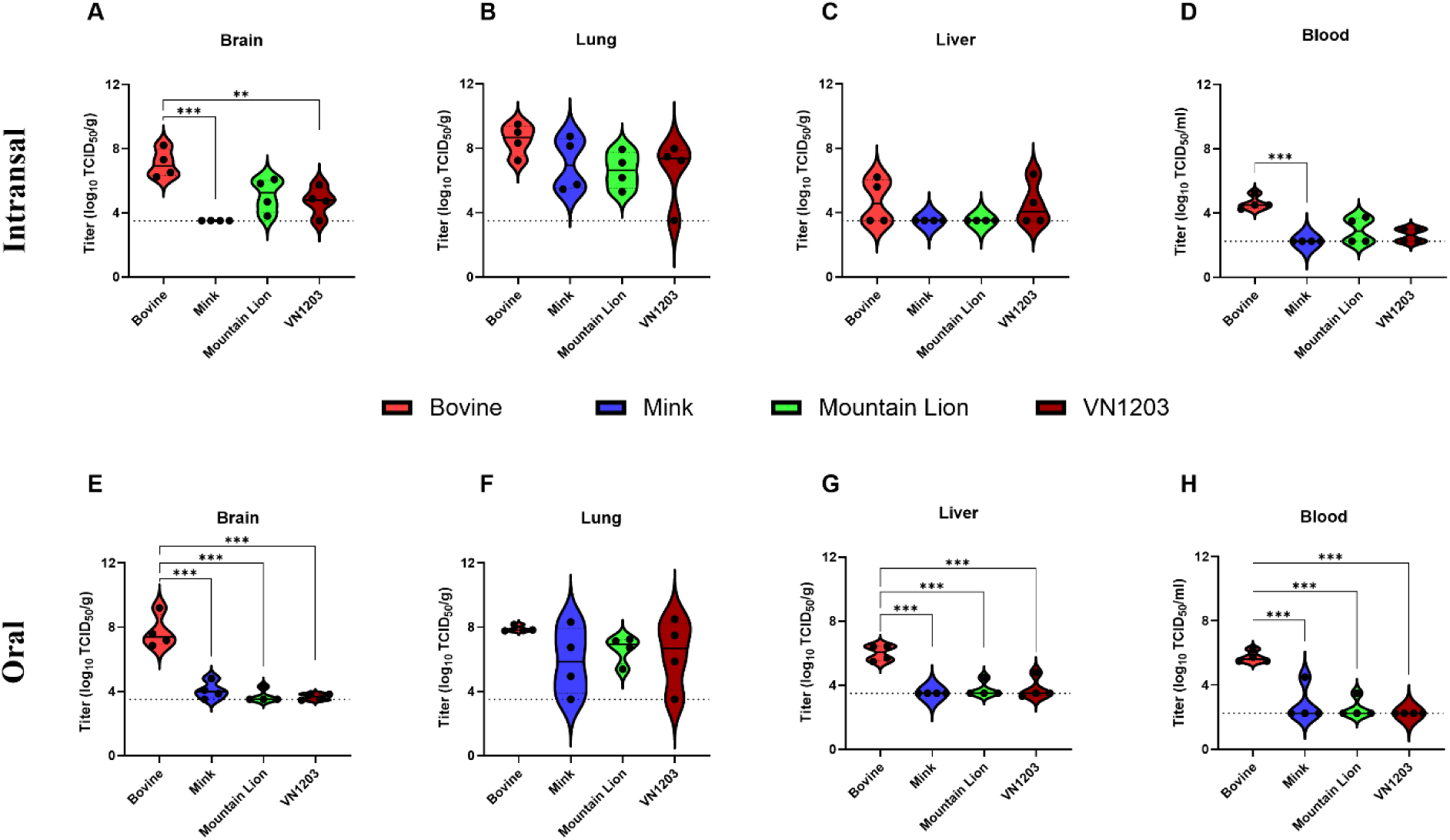
Infectious virus titers in organs following HPAI A(H5N1) infection. Six-week-old C57BL/6J mice were inoculated with 10^5^ TCID50 of A/bovine/OH/B24OSU-342/2024, A/mountain lion/MT/1/2024), A/mink/Spain/3691-8_22VIR10586-10/2022, or A/Vietnam/1203/2004. On day 4 post-inoculation, 4 animals from each group were euthanized determine virus titers in organs (lung, liver, brain) and blood loads. (A-D) Intranasally inoculated mice. (E-H) Orally inoculated mice. Dashed line indicates limit of detection. Statistical analyses were performed using one-way ANOVA with Tukey’s multiple comparison. ns p > 0.05, *p < 0.05, **p < 0.01, ****p < 0.0001.

### Bovine isolate caused virus replication in brain tissue

To determine specific differences in pathogenesis by the HPAI A(H5N1) isolates in C57BL/6J mice, we next assessed histologic lesions in brain and lung samples collected on day 4 PI. Regardless of route of inoculation or isolate, no significant histopathologic changes were observed in evaluated brain sections (**Fig. 3A,E,I,M; Fig 4A,E,I,M**). Abundant nucleoprotein antigen was observed in neurons, glial cells and occasional ependymal cells in mice intranasally and orally inoculated with the bovine isolate in the absence of tissue inflammation (**Fig. 3B**; **Fig. 4B**). No virus antigen was observed in any brain cell type of mice intranasally and orally inoculated with the mink and VN1203 isolates (**Fig. 3F,N**; **Fig. 4F,N**). For the mountain lion isolate, rare and minimal virus antigen was observed in the brain of two animals (one of four each intranasally and orally) (**Fig. 3J**; **Fig. 4J**) but the majority of animals inoculated with the mountain lion isolate did not have observable antigen in the brain. Overall, virus antigen was detected in the absence of visible evidence of inflammation in brain sections of animals infected with the bovine isolate.

**Figure 3:**
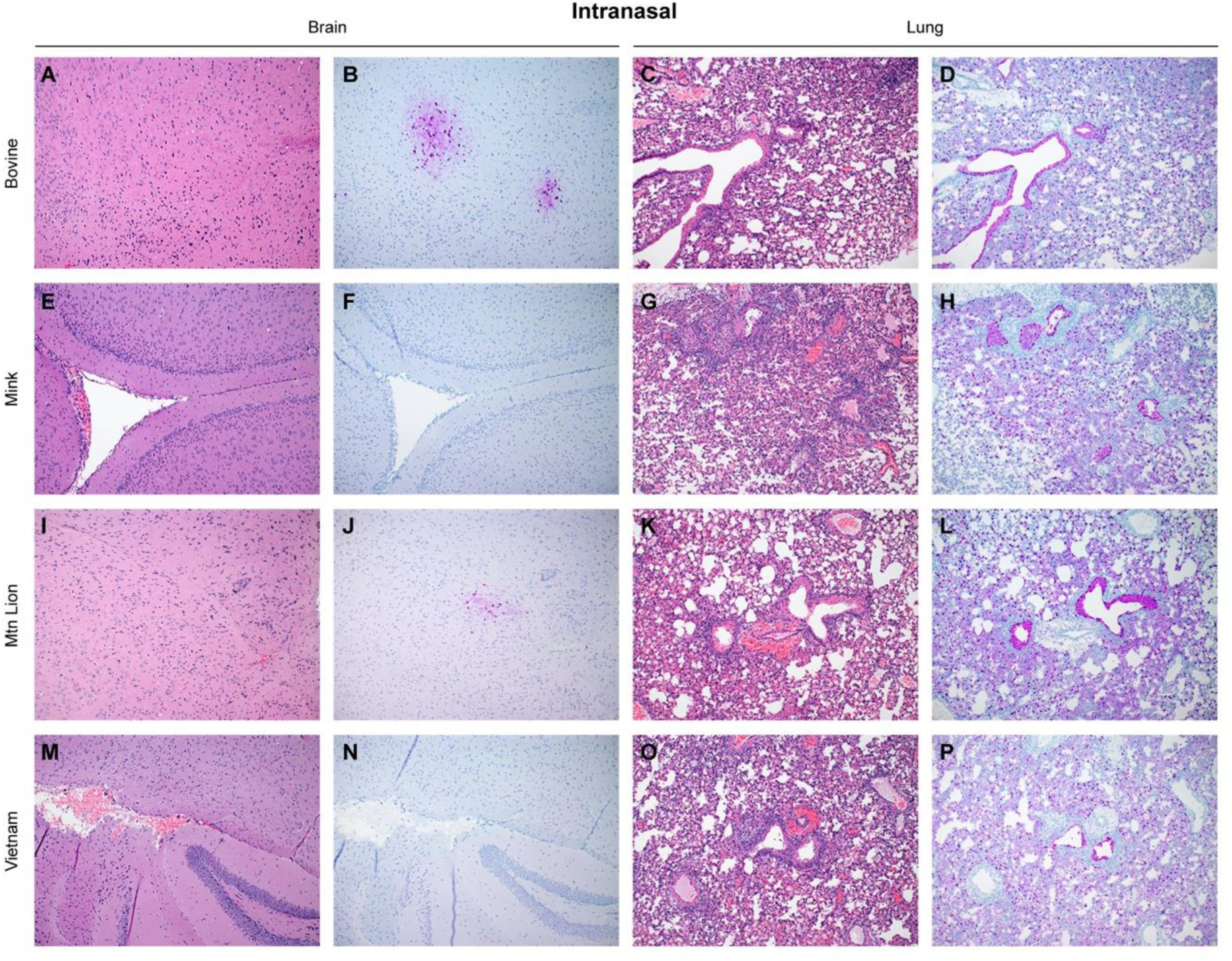
Histologic lesions and antigen expression following HPAI A(H5N1) intranasal inoculation. Six-week-old C57BL/6J mice were inoculated intranasally with 10^5^ TCID50 of A/bovine/OH/B24OSU-342/2024, A/mountain lion/MT/1/2024), A/mink/Spain/3691- 8_22VIR10586-10/2022, or A/Vietnam/1203/2004. A subset of animals (n = 4) was euthanized at day 4 post-inoculation for histopathologic evaluation. Section from immunoreactive mice with histopathologic changes shown. A,E,I,M – hematoxylin & eosin stain of brain sections; C,G,K,O - hematoxylin & eosin stain of lung sections; B,F,J,N – immunohistochemistry staining of brain sections for Influenza A virus nucleoprotein antigen; D,H,L,P - immunohistochemistry staining of lung sections for Influenza A virus nucleoprotein.

**Figure 4:**
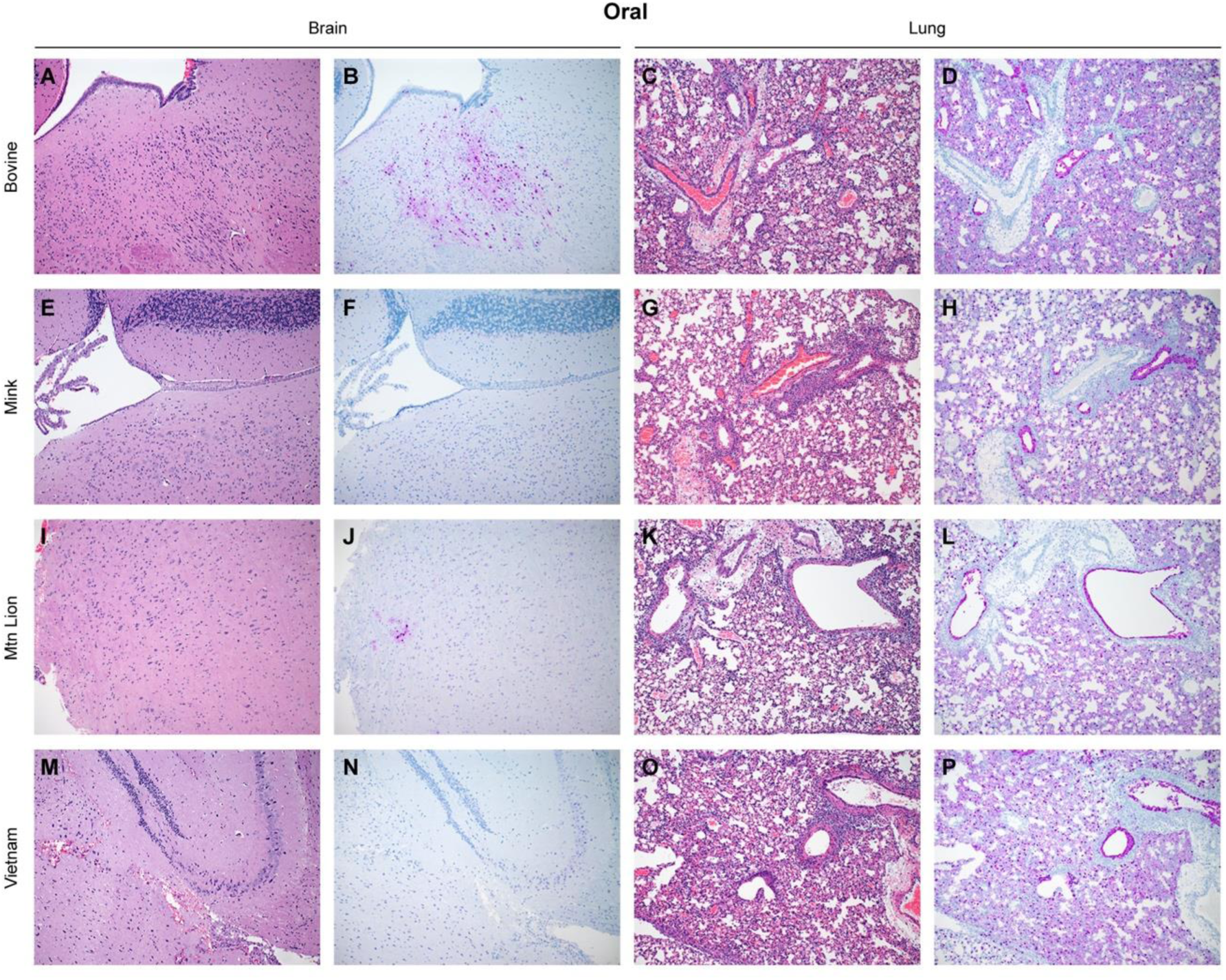
Histologic lesions and antigen expression following HPAI A(H5N1) oral inoculation. Six-week-old C57BL/6J mice were inoculated orally with 10^5^ TCID50 of A/bovine/OH/B24OSU-342/2024, A/mountain lion/MT/1/2024), A/mink/Spain/3691- 8_22VIR10586-10/2022, or A/Vietnam/1203/2004. A subset of animals from each group (n = 4) was euthanized at day 4 post-infection for histopathologic evaluation. Section from immunoreactive mice with histopathologic changes shown. A,E,I,M – hematoxylin & eosin stain of brain sections; C,G,K,O - hematoxylin & eosin stain of lung sections; B,F,J,N – immunohistochemistry staining of brain sections for Influenza A virus nucleoprotein antigen; D,H,L,P - immunohistochemistry staining of lung sections for Influenza A virus nucleoprotein.

Histologic changes were observed in lungs from both intranasally and orally inoculated mice from each isolate (**Fig. 3C,G,K,O; Fig. 4C,G,K,O**). Mild to severe necrotizing bronchiolitis with minimal to moderate interstitial pneumonia was observed in all mice inoculated with the bovine isolate (**Fig. 3C;4C**). Similarly, mice intranasally inoculated with the mountain lion isolate also showed moderate to severe bronchiolitis and interstitial pneumonia (**Fig. 3K**) whereas disease in the orally inoculated mice was less severe with moderate necrotizing bronchiolitis and interstitial pneumonia observed (**Fig. 4K**). In contrast, an inconsistent histopathologic picture was observed with the mink isolate. Lungs from mice intranasally inoculated with the mink isolate exhibited minimal to severe bronchiolitis with moderate interstitial pneumonia (**Fig. 3G**). Similarly, in the orally inoculated mice, there was a large range of observable pathology from no significant histopathologic changes to one mouse having severe necrotizing bronchiolitis with moderate interstitial pneumonia for oral inoculation (**Fig. 4G**).

This histologic picture suggests limited pathology on day 4 PI and matches with slower disease progression observed in mice inoculated with the mink isolate. Severe bronchiolitis and moderate interstitial pneumonia was observed in mice intranasally inoculated with the VN1203 isolate (**Fig. 3O**) while two of the four orally inoculated mice presented with severe bronchiolitis with mild to moderate interstitial pneumonia (**Fig. 4O**) and the remaining mice showing no significant histopathologic lesions.

Immunohistochemical analysis closely mirrored histopathologic observations in the lung. Nucleoprotein antigen was detected in bronchiolar epithelium, type I and II pneumocytes and alveolar macrophages of all mice inoculated with the bovine isolate (**Fig. 3D**; **Fig. 4D**).

Similarly, virus antigen was observed in every evaluated lung section of mice intranasally and orally inoculated with the mountain lion isolate (**Fig. 3L**; **Fig. 4L**). For two of the four mice intranasally and one out of four mice orally inoculated with the mink isolate, virus antigen was detected in bronchiolar epithelium, type I and II pneumocytes and alveolar macrophages with limited virus antigen detected in bronchiolar epithelial cells (**Fig. 3H**; **Fig.4H**). Virus antigen detection mimicked lesion severity as the most severe changes correlated to the most immunoreactive cell types and mice that had not histopathologic changes exhibited no viral nucleoprotein immunoreactivity. Similarly, virus antigen was detected in all segments of the conducting airways in mice intranasally infected with the VN1203 isolate (**Fig. 3P**) whereas in orally inoculated mice virus antigen was identified in two of the four evaluate mice (**Fig. 4P**). Overall, there was a strong correlation between histopathologic changes and virus antigen detection in the lung supporting high viral lung loads and respiratory disease in infected mice.

### Bovine isolate induced pro-inflammatory cytokine profile in the brain

Pathogenesis following inoculation with HPAI A(H5N1) strains has been associated with increased production of cytokines, chemokines and IFNs in humans ^16^. We next determined whether the observed differences in disease course by the different HPAI A(H5N1) isolates correlated with cytokine dysregulation in our model. Since C57BL/6J mice are known to mount a predominantly Th1 response ^17^, we evaluated the cytokines in lungs and brains of inoculated mice with a focus on the Th1 response. In orally inoculated mice, significant reduction in IL-1β and GM-CSF cytokine levels was observed with the bovine isolate compared to the mink isolate in lungs (**Fig. S1**) whereas IL-6 was significantly reduced in both brains and lungs from VN1203 inoculated mice compared to bovine inoculated mice (**Fig. S1; Fig. S2**). No significant differences (between the bovine isolate and each of mink, mountain lion, VN1203 isolates) in inflammatory cytokine levels were observed in lungs from intranasally inoculated mice (**Fig. S3**). Significantly higher pro-inflammatory IL-1α, IL-1β, IL-2 and GM-CSF cytokine levels were observed in brain tissues from animals inoculated intranasally with the bovine isolate compared to mink, mountain lion and VN1203 isolates (**Fig. 5**). The results are consistent with the detection of infectious virus and viral antigen in the brains of C57BL/6J mice infected with the bovine isolate.

**Figure 5:**
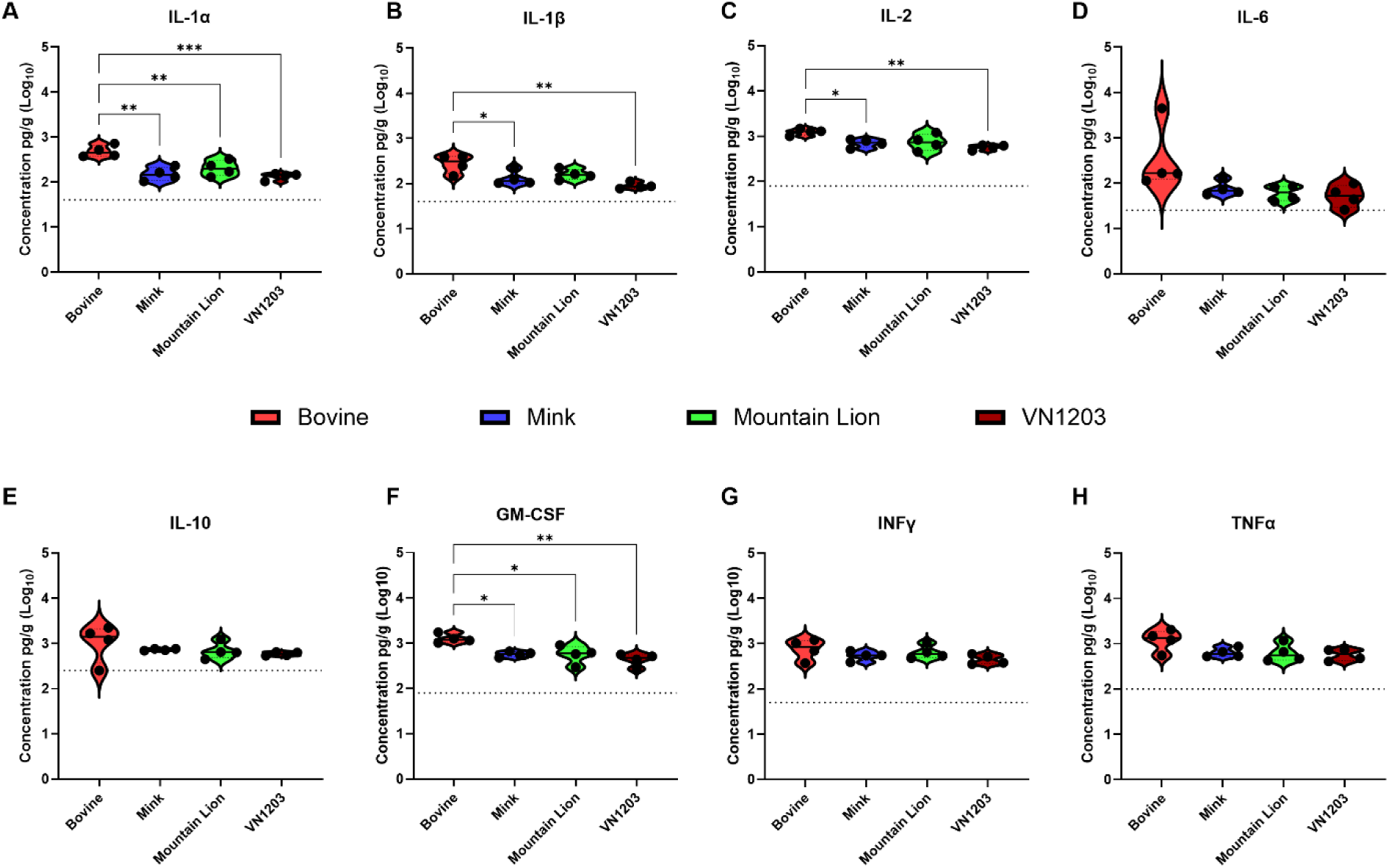
Brain cytokine expression following HPAI A(H5N1) infection. Six-week-old C57BL/6J mice (n = 10 per group) were inoculated with 10^5^ TCID50 of A/bovine/OH/B24OSU- 342/2024, A/mountain lion/MT/1/2024), A/mink/Spain/3691-8_22VIR10586-10/2022, or A/Vietnam/1203/2004. A subset of animals from each group (n = 4) was euthanized at day 4 post infection for cytokine analysis. (A-D) Intranasally inoculated mice. (E-H) Orally inoculated mice. Dashed line indicates limit of detection. Statistical analyses were performed using one-way ANOVA with Tukey’s multiple comparison. ns p > 0.05, *p < 0.05, **p < 0.01, ****p < 0.0001.

### Bovine isolate caused more rapid respiratory disease in BALB/c mice

To reinforce our initial findings in C57BL/6J mice we utilized the same experimental set up with groups of BALB/c mice (n = 10/group). BALB/c mice were intranasally or orally inoculated with 10^5^ TCID50 of A/bovine/OH/B24OSU-342/2024, A/mountain lion/MT/1/2024, A/mink/Spain/3691- 8_22VIR10586-10/2022/ and A/Vietnam/1203/04, and animals were weighed daily and monitored for signs of illness. All inoculated mice developed weight loss (**Fig. S4A,B**), ruffled fur, and hunched posture followed by rapid breathing but no obvious neurologic signs. BALB/c mice inoculated with the bovine strain all succumbed a day earlier compared to inoculated C57BL/6J mice. All mice inoculated intranasally with the different isolates succumbed to infection except for a single mouse in the mink group (**Fig. S4C**). Like C57BL/6J mice, oral inoculations with the mink, mountain lion and VN1203 isolates did not result in uniform lethality (**Fig. S4D**). For comparison of viral load and disease, BALB/c mice were euthanized on day 2 PI when they had reached a similar disease stage as the inoculated C57BL/6 mice (**Fig. S5A-H**). In contrast to C57BL/6J mice, virus brain titers in mice inoculated with all isolates were negligible and in line with the respiratory disease observed in BALB/c mice (**Fig. S5A,E**). All virus isolates replicated at similar levels in the BALB/c mice with high titers in the lungs (**Fig. S5B,F**) regardless of inoculation route. Only the bovine isolate caused significant viremia and infectious virus was found in blood samples of 3/4 intranasally and in 4/4 orally infected BALB/c mice (**Fig S5D,H**). Cytokine induction by the bovine isolate was not significantly different compared to mink, mountain lion and VN1203 isolates regardless of route of infection (**Fig. S6; Fig. S7; Fig. S8; Fig S9**). Overall, all HPAI A(H5N1) isolates used here caused respiratory infection in BALB/c mice lacking evidence of neurovirulence. Virus replication revealed similar organ titers for all isolates with highest titers in lungs. Disease progression in mice infected with the bovine isolate was more advanced than with the mink, mountain lion and VN1203 isolates.

## DISCUSSION

Recent increases in cases of HPAI A(H5N1) clade 2.3.4.4b virus infections in mammals including humans is concerning for animal and public health^18^. Human cases in the United States remain mild and limited in number but continued circulation with incidental spillover into mammalian populations could confer mutations resulting in more efficient transmission and increased virulence in mammals including humans.

Our mouse studies have demonstrated increased virulence of a recent bovine isolate over the reference VN1203 and two distinct recent HPAI A(H5N1) isolates (mink and mountain lion) in two mouse strains. We show that the bovine isolate caused systemic infection with increased capacity for neuroinvasion and neurovirulence a trade that is not unfamiliar with HPAI A(H5N1) clade 2.3.4.4b viruses ^4,14,19^. Interestingly, systemic infection was not obvious with the mink, mountain lion and VN1203 isolates showing only insignificant and inconsistent viremia in individual infected animals. The disease course observed in C57BL/6J mice inoculated with the bovine isolate is a consequence of combined neurological and respiratory disease. High viral organ loads, antigen staining and an elevated pro-inflammatory cytokine profile in brain tissue further support neurological disease manifestation associated with the bovine isolate. There was some evidence of neurological involvement with the mountain lion isolate with a minority of mice showing neurologic disease signs and antigen staining in the brain. Noteworthy, the mountain lion and bovine isolates are genetically more closely related.

The mechanism of neuroinvasion of the bovine isolate remains undetermined but the presence of viremia may favor hematogenous spread via the blood-brain-barrier. Indeed, bloodborne spread of an H5N1 isolate and a mouse adapted variant of H7N7 has been documented ^20,21^.

Neuroinvasion through cranial nerves, which seems plausible for the intranasal route of infection, also remains a viable option, and both neuroinvasion through the blood-brain-barrier and cranial nerves have been demonstrated in ferrets infected with H5N1 ^22^. Further investigations are needed to understand virus replication and spread for the oral route of infection, although submucosal and myenteric plexi in the intestine could serve as sites of virus entry. We cannot, however, exclude that the mice may have been unintendedly co-exposed via the respiratory tract during oral inoculation.

Interestingly, neurologic disease and infectivity in brain tissue was not observed with BALB/c mice infected with the bovine isolate despite systemic infection. This could be a consequence of advanced virus replication in BALB/c over C57BL/6J mice leading to fatal respiratory disease before neurovirulence may have occurred. A recent paper also did not describe neurologic disease in BALB/c mice infected with the bovine isolate supporting our result here ^23^.

Conversely, there may be a distinct organ tropism and disease progression associated with the bovine isolate in the two mouse strains. BAB/c and C57BL/6J mice differ in their immune responses with C57BL/6J mice demonstrating a biased Th1 and M-1 response and BALB/c mice showing a dominant Th2 and M-2 response^17,24^. These differences in immune responses have been discussed for differences in susceptibility of these mice to the 2009 pandemic H1N1 influenza virus and a human H5N1 isolate ^25^. Currently, it remains unclear whether differences in baseline immune responses influence disease progression and pathology of current H5N1 isolates.

High viral load and viral antigen staining in brain tissue indicated replication of the bovine isolate in the central nervous system. The absence of histological lesions, however, was intriguing as infected mice developed neurological signs. Given the lack of detectable inflammatory infiltrates and similar lack of neuronal necrosis, neuronal metabolic disturbances are deemed the most probable explanation for the described clinical signs. The proinflammatory phenotype found in brain tissue may support this notion. Necrosis and inflammation in brain tissue notably in the cerebrum has been reported before in H5N1 infected mammals ^4,20^ which we did not observe here.

Certain molecular markers/motifs have been reported to influence the replicative capacity, virulence, and transmission of influenza A viruses in both poultry and mammals ^26,27^. All recent HPAI H5N1 isolates used here still possess the full-length neuraminidase (NA) stalk region, which is lacking in the VN1203 isolate. Deletion of this stalk region has been associated with increased virulence in avian species and mice ^28–31^. Similarly, a 5-amino acid deletion in the nonstructural protein NS1 linker region connecting the RNA binding domain with the effector domain has been reported to confer increased viral replication potential in chickens and mice ^32–34^. Again, this deletion is not present in the three recent isolates in contrast to VN1203 indicating that other mutations may be equally or more important for virulence in mice. Furthermore, the E627K substitution in PB2, one of the polymerase complex proteins, of avian-origin influenza viruses is thought to support more efficient replication in mammalian species including mice ^35,36^. In contrast to VN1203, none of the recent HPAI A(H5N1) isolates used in this study possesses this substitution despite similar or higher virulence in mice. On the other hand, all HPAI A(H5N1) isolates evaluated in the study harbor mutations in the matrix protein M1 (such as 30D, 43M, and 215A) ^26,37,38^ and NS1 (such as 42S, 103F, and 106M)^39^ known to increase virulence in mice. Future studies need to perform closer analyses of all the mutational differences to better understand and define molecular signatures that explain the increased virulence and altered organ tropism of the bovine isolate in mammalian species.

Our study defined oral inoculations as a potent exposure route for HPAI A(H5N1) clade 2.3.4.4b viruses. This offers a plausible explanation for reported infections in wild and domesticated carnivores with HPAI A(H5N1) viruses that may get infected through hunting and scavenging as may be the case for the mountain lion from which one of the viruses used here was isolated. In addition, oral exposure may be a viable infection route for other mammals, such as cats and humans, when consuming raw milk from infected dairy cows. Oral exposure could also be a route supporting cattle-to-cattle transmission. More studies would be necessary to determine the risk of infection by the oral-gastric route.

In conclusion, the recent bovine isolate showed high virulence for mice when inoculated by the intranasal and oral routes. C57BL/6J and BALLB/c mice provide excellent in vivo screening models for efficacy testing of antiviral, therapeutic and vaccine candidates against infections with these emerging HPAI A(H5N1) viruses. The C57BL/6J model allows studies on mechanisms of neuroinvasion and neurovirulence of the emerging HPAI A(H5N1) viruses. Both mouse models may be used to gain deeper insight into infection by the oral-gastric route, an exposure route of importance for cow-to-human as well as bird/poultry-to-carnivore transmission of emerging HPAI A(H5N1) isolates. Thus, the mouse models will be extremely beneficial for animal and public health responses to the current HPAI A(H5N1) outbreak.

## METHODS

### Biosafety and ethics

All infectious work was conducted at maximum biosafety level following operating procedures approved by the Institutional Biosafety Committee of the Rocky Mountain Laboratories, Division of Intramural Research, National Institute of Allergy and Infectious Diseases, National Institutes of Health (Hamilton, MT, USA). Animal experiments were approved by the Rocky Mountain Laboratories Animal Care and Use Committee (ACUC) in an AALAC accredited facility. Sample inactivation followed established protocols approved by the RML Institutional Biosafety Committee.

### Viruses and cells

The highly pathogenic H5N1 virus strains used in this study include A/Vietnam/1203/2004, A/bovine/OH/B24OSU-342/2024, A/mink/Spain/3691-8_22VIR10586- 10/2022 ^9^ and A/mountain lion/MT/1/2024 ^10^. Virus growth and titration was performed on Madin-Darby canine kidney (MDCK) cells in Minimal Essential Media (MEM) (Sigma-Aldrich, St. Louis, MO) containing 4µg/ml trypsin, 2mM L-glutamine, 50U/mL penicillin, 50µg/ml streptomycin, 1% NEAA, 20mM HEPES (all Thermo Fisher Scientific, Waltham, MA). MDCK cells were maintained in MEM supplemented with 10% fetal bovine serum (FBS) (Wisent Inc., St. Bruno, Canada), 2mM L-glutamine, 50U/mL penicillin, 50μg/ml streptomycin, 1% non-essential amino acids (NEAA) and 20mM HEPES.

### Mouse inoculations

Six-week-old C57BL/6J and BALB/c mice were purchased from Jackson laboratories and randomly assigned to study groups. Ten mice per virus group were anaesthetized with isoflurane and inoculated with 10^5^ TCID50 of A/bovine/OH/B24OSU- 342/2024, A/mink/Spain/3691-8_22VIR10586-10/2022, A/mountain lion/MT/1/2024) and A/Vietnam/1203/04 intranasally or orally in volumes of 50µl and 250µl, respectively. All animals were weighed daily and monitored for clinical signs of disease. Four animals per inoculation group were euthanized on day 2 post-infection (BALB/c mice) or day 4 post-infection (C57BL/6J mice) at which time blood and tissues (brain, lung, liver) were collected for virus titration and histopathology. Six mice per inoculation group were monitored for survival until day 28 post inoculation, anesthetized, bled and humanely euthanized.

### Virus titration

Infectious virus was quantified on MDCK cells. MDCK cells were seeded in 96- well plates in MEM supplemented with 10% FBS, 2mM L-glutamine, 50U/mL penicillin, 50μg/ml streptomycin, 1% NEAA and 20mM HEPES. Mouse tissue specimens in tubes containing steel beads and 1 mL infection media (MEM containing 4μg/ml trypsin, 2mM L-glutamine, 50U/mL penicillin, 50μg/ml streptomycin, 1x NEAA and 20mM HEPES) were homogenized using the TissueLyser II (Qiagen). Blood samples were initially diluted (1:10) in phosphate-buffered saline without Ca2^+^/Mg2^+^. Subsequent ten-fold serial dilutions were performed in media. A 100 μL aliquot of diluted sample was applied on MDCK cells and incubated for 3 days at 37°C. Subsequently, 75µl of 0.33% Turkey erythrocytes were added to 25 µl aliquot of the supernatant from infected MDCK cells in 96 well plates and incubated for 1 hour at 37°C before hemagglutination was read. TCID50 was calculated using the Reed & Muench method ^40^.

### Cytokine analysis

Brain and lung tissue samples were thawed and homogenized in 1ml of phosphate buffered saline by using a TissueLyser II at 30-Hz oscillation frequency for 5 min. Homogenates were centrifuged (10,000 × *g* for 5 min) to remove debris and irradiated according to approved procedures to inactivate HPAI H5N1 virus ^41^. Cleared supernatants were used for cytokine quantification using the Bio-Plex Pro^TM^ Mouse Cytokine 23-plex Assay (Biorad) according to the manufacturer’s instructions.

### Histopathology and Immunohistochemistry

Tissues were fixed in 10% Neutral Buffered Formalin x2 changes, for a minimum of 7 days. Tissues were placed in cassettes and processed with a Sakura VIP-6 Tissue Tek, on a 12-hour automated schedule, using a graded series of ethanol, xylene, and ParaPlast Extra. Embedded tissues are sectioned at 5um and dried overnight at 42 degrees Celsius prior to staining. Immunoreactivity was detected using Millipore Sigma Anti-Influenza A nucleoprotein antibody (Cat.#ABF1820-25UL) at a 1:12,000 dilution. Roche Tissue Diagnostics DISCOVERY Omnimap anti-rabbit HRP (#760-4311) was used as a secondary antibody. For negative controls, replicate sections from each control block were stained in parallel following an identical protocol, with the primary antibody replaced by Vector Laboratories rabbit IgG (#I-1000-5) at a 1:2500 dilution. The tissues were stained using the DISCOVERY ULTRA automated stainer (Ventana Medical Systems) with a Roche Tissue Diagnostics DISCOVERY purple kit (#760-229).

### Amino acid sequence analysis

Amino acid identity of all the proteins of HPAI H5N1 isolate VN1203, Mink, Mountain Lion, and Bovine were analyzed using Clustal Omega, a multiple sequence alignment tool^42^.

### Statistical analysis

Statistical analysis was performed using GraphPad Prism version 10.3.0 for Windows (GraphPad Software, San Diego, CA, USA, www.graphpad.com, 4 July 2024).

Differences between mouse groups were compared using one-way ANOVA with Turkey’s multiple comparison. Survival curves were compared by using the log-rank Mantel–Cox test. Statistical significance was set at p < 0.05.

## ACKNOWLEDGEMENTS

We thank Richard Webby from St. Judes Children hospital and Andrew Bowman from Ohio State university for providing the bovine isolate. The mink isolate was provided by Isabella Monne & Calogero Terregino, Istituto Zooprofilattico Sperimentale delle Venezie; Monserrat Agüero, Laboratorio Central de Veterinaria, Spain, Algete, Madrid; Gustavo del Real, Centro National de biotechnologica, Madrid Spain. We are grateful to Brandi Williamson and Tessa Luttermann (Division of Intramural Research, NIAID) for support with virus growth and training. The authors also thank the Rocky Mountain Veterinary Branch, NIAID for animal care and histopathology service as well as the High Containment support staff for general biocontainment service.

This work was supported by the Intramural Research Program of the NIAID.

## Author Contributions

TT, KR and HF designed the study. VJM, EW and KCY acquired virus isolates, grew virus stock and confirmed virus sequences. TT, KR, MV, KMW, ML, AO, NM, CC performed experiments and data analysis. TT, MV, KR and HF wrote the manuscript. HF obtained funding. All authors reviewed and contributed to preparation of the final manuscript.

