## Supplementary material for "Recent Bovine HPAI H5N1 Isolate Is Highly Virulent For Mice, Rapidly Causing Acute Pulmonary And Neurologic Disease"

### Slide 1
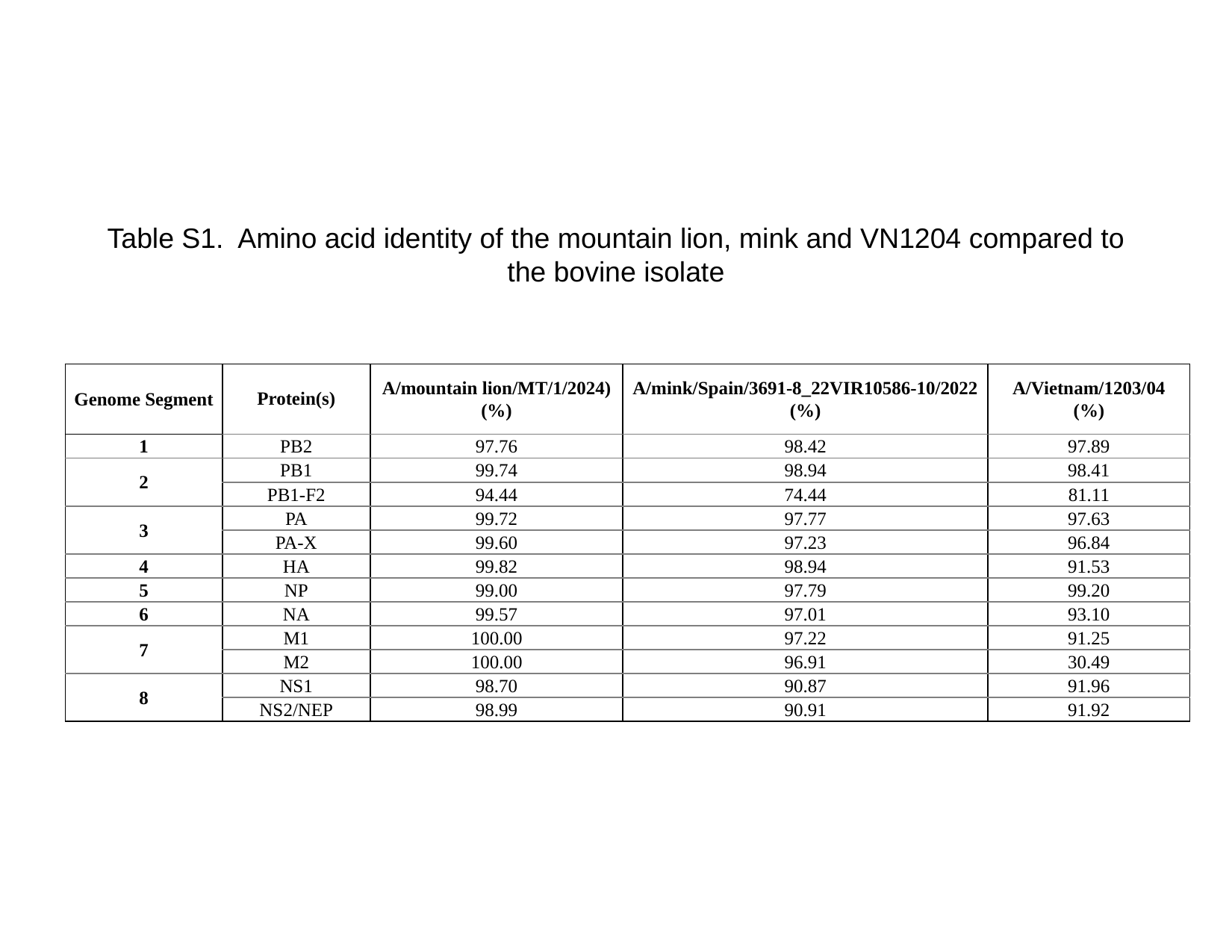

Table S1. Amino acid identity of the mountain lion, mink and VN1204 compared to the bovine isolate
| Genome Segment | Protein(s) | A/mountain lion/MT/1/2024) (%) | A/mink/Spain/3691-8\_22VIR10586-10/2022 (%) | A/Vietnam/1203/04 (%) |
| --- | --- | --- | --- | --- |
| 1 | PB2 | 97.76 | 98.42 | 97.89 |
| 2 | PB1 | 99.74 | 98.94 | 98.41 |
| | PB1-F2 | 94.44 | 74.44 | 81.11 |
| 3 | PA | 99.72 | 97.77 | 97.63 |
| | PA-X | 99.60 | 97.23 | 96.84 |
| 4 | HA | 99.82 | 98.94 | 91.53 |
| 5 | NP | 99.00 | 97.79 | 99.20 |
| 6 | NA | 99.57 | 97.01 | 93.10 |
| 7 | M1 | 100.00 | 97.22 | 91.25 |
| | M2 | 100.00 | 96.91 | 30.49 |
| 8 | NS1 | 98.70 | 90.87 | 91.96 |
| | NS2/NEP | 98.99 | 90.91 | 91.92 |

### Slide 2
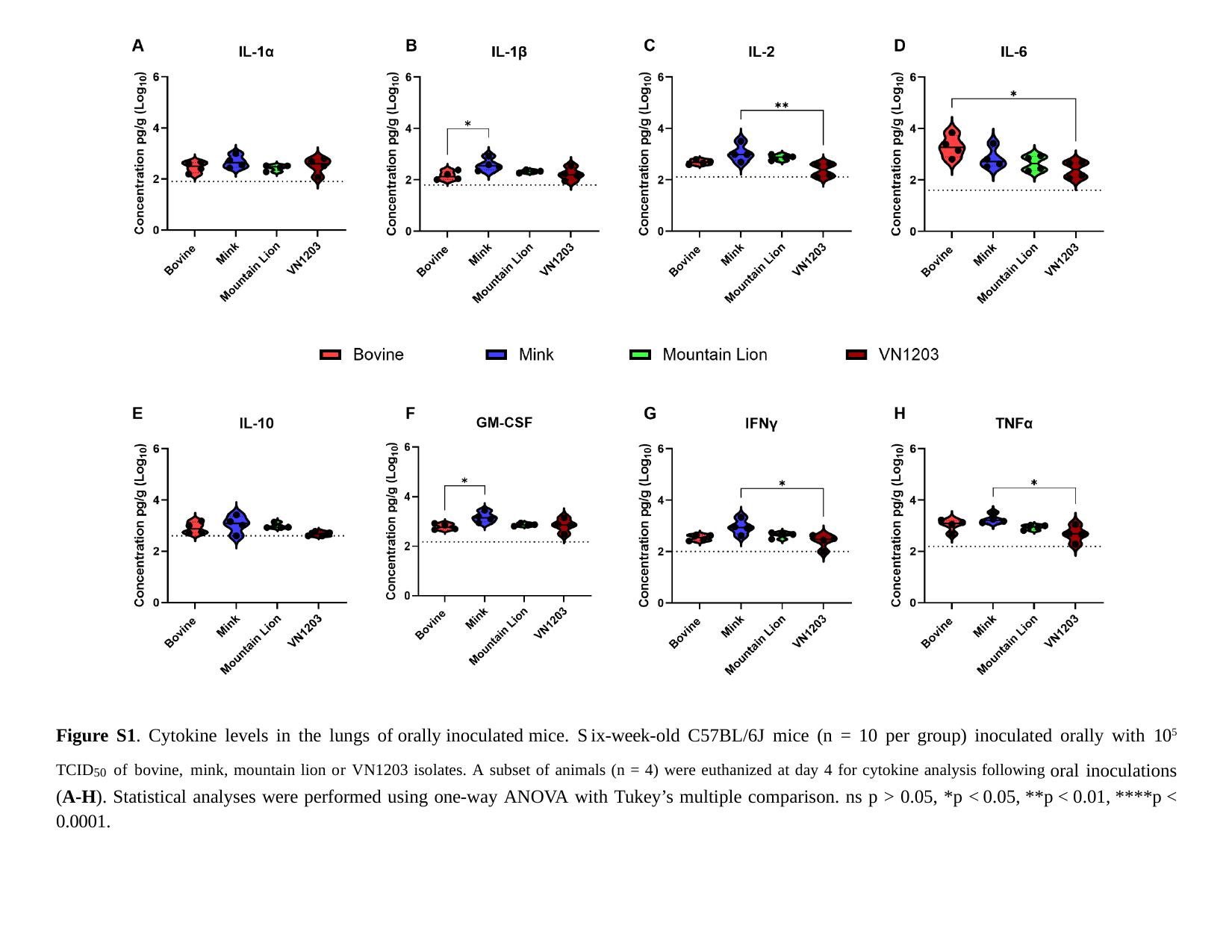

Figure S1. Cytokine levels in the lungs of orally inoculated mice. Six-week-old C57BL/6J mice (n = 10 per group) inoculated orally with 10^5^ ^TCID^50 ^of^ ^bovine,^ ^mink,^ ^mountain^ ^lion^ ^or^ ^VN1203^ ^isolates.^ ^A^ ^subset^ ^of^ ^animals^ ^(n^ ^=^ ^4)^ ^were^ ^euthanized^ ^at^ ^day^ ^4^ ^for^ ^cytokine^ ^analysis^ ^following^ oral inoculations (A-H). Statistical analyses were performed using one-way ANOVA with Tukey’s multiple comparison. ns p > 0.05, *p < 0.05, **p < 0.01, ****p < 0.0001.

### Slide 3
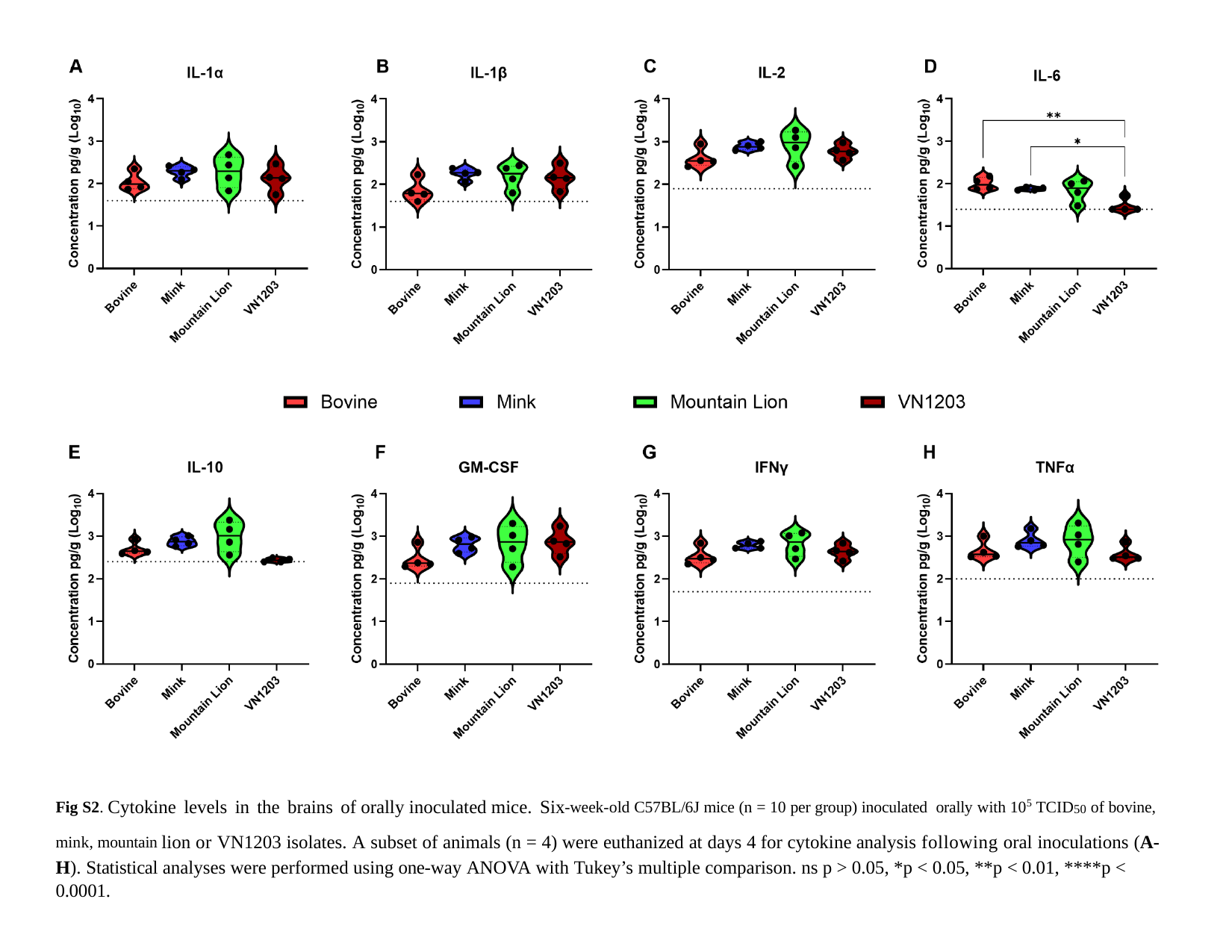

^Fig^ ^S2.^ Cytokine levels in the brains of orally inoculated mice. Six^-week-old^ ^C57BL/6J^ ^mice^ ^(n^ ^=^ ^10^ ^per^ ^group)^ ^inoculated^ ^orally^ ^with^ ^105^ ^TCID^50 ^of^ ^bovine,^ ^mink,^ ^mountain^ lion or VN1203 isolates. A subset of animals (n = 4) were euthanized at days 4 for cytokine analysis following oral inoculations (A-H). Statistical analyses were performed using one-way ANOVA with Tukey’s multiple comparison. ns p > 0.05, *p < 0.05, **p < 0.01, ****p < 0.0001.

### Slide 4
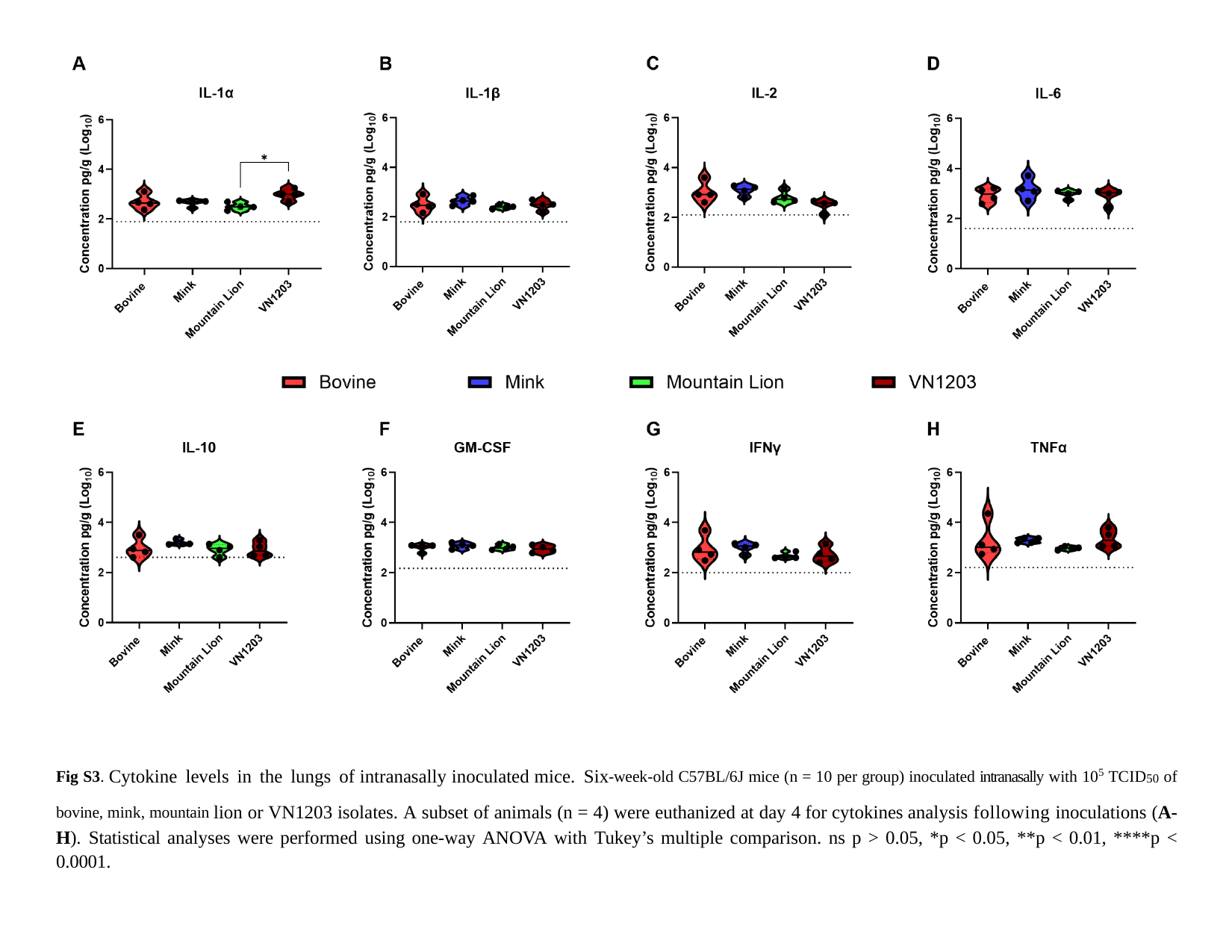

^Fig^ ^S3.^ Cytokine levels in the lungs of intranasally inoculated mice. Six^-week-old^ ^C57BL/6J^ ^mice^ ^(n^ ^=^ ^10^ ^per^ ^group)^ ^inoculated^ ^intranasally^ ^with^ ^105^ ^TCID^50 ^of^ ^bovine,^ ^mink,^ ^mountain^ lion or VN1203 isolates. A subset of animals (n = 4) were euthanized at day 4 for cytokines analysis following inoculations (A-H). Statistical analyses were performed using one-way ANOVA with Tukey’s multiple comparison. ns p > 0.05, *p < 0.05, **p < 0.01, ****p < 0.0001.

### Slide 5
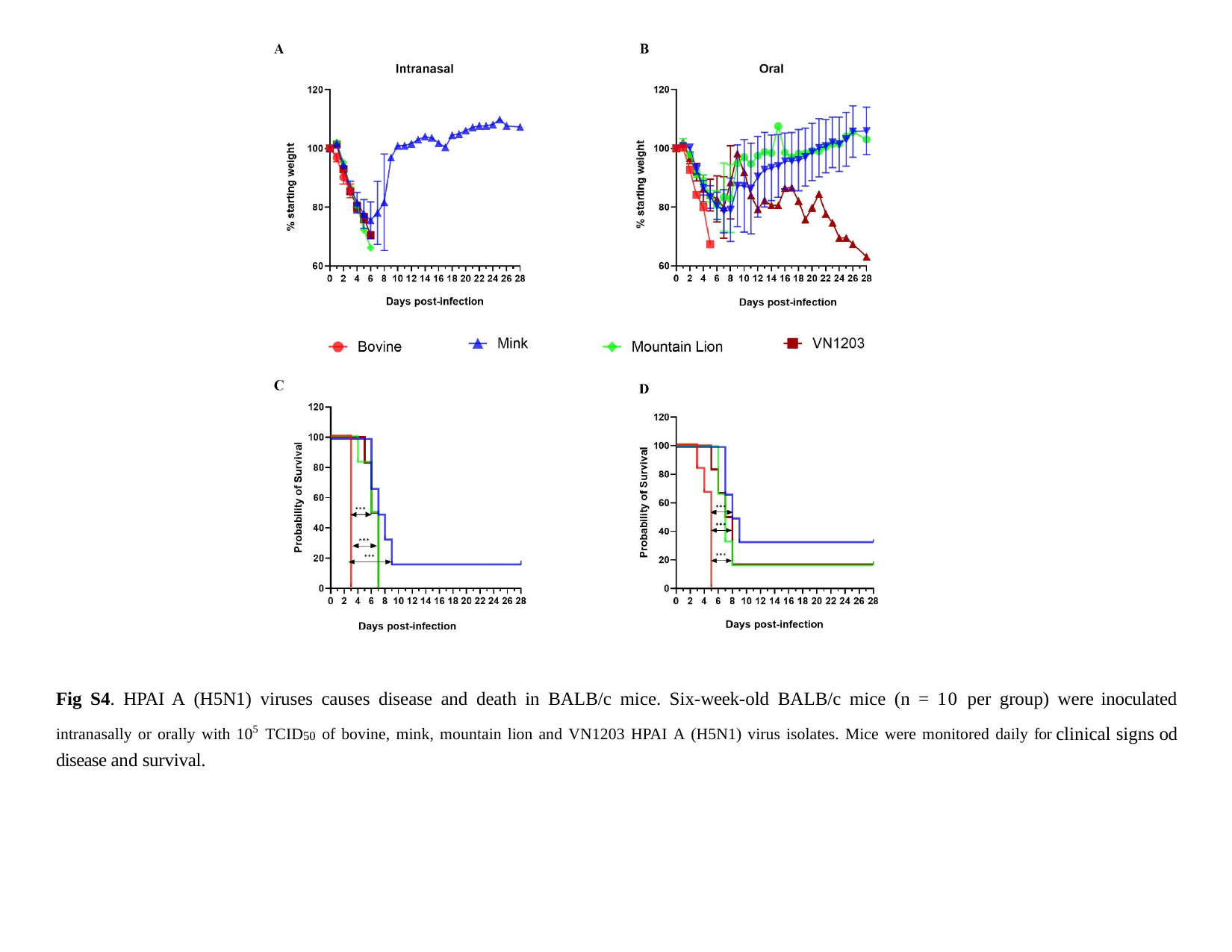

Fig S4. HPAI A (H5N1) viruses causes disease and death in BALB/c mice. Six-week-old BALB/c mice (n = 10 per group) were inoculated ^intranasally^ ^or^ ^orally^ ^with^ ^105^ ^TCID^50 ^of^ ^bovine,^ ^mink,^ ^mountain^ ^lion^ ^and^ ^VN1203^ ^HPAI^ ^A^ ^(H5N1)^ ^virus^ ^isolates.^ ^Mice^ ^were^ ^monitored^ ^daily^ ^for^ clinical signs od disease and survival.

### Slide 6
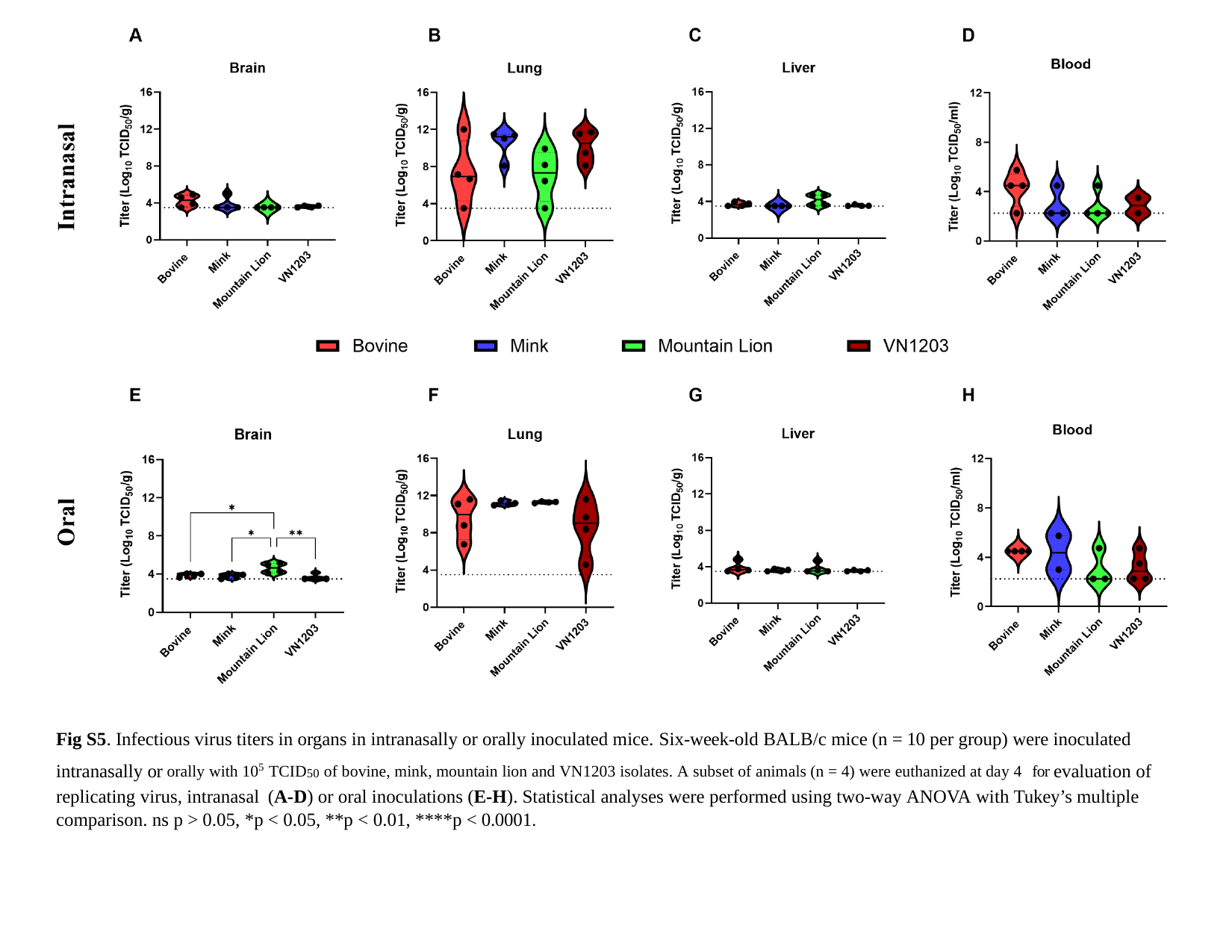

Fig S5. Infectious virus titers in organs in intranasally or orally inoculated mice. Six-week-old BALB/c mice (n = 10 per group) were inoculated intranasally or ^orally^ ^with^ ^105^ ^TCID^50 ^of^ ^bovine,^ ^mink,^ ^mountain^ ^lion^ ^and^ ^VN1203^ ^isolates.^ ^A^ ^subset^ ^of^ ^animals^ ^(n^ ^=^ ^4)^ ^were^ ^euthanized^ ^at^ ^day^ ^4^ ^for^ evaluation of replicating virus, intranasal (A-D) or oral inoculations (E-H). Statistical analyses were performed using two-way ANOVA with Tukey’s multiple comparison. ns p > 0.05, *p < 0.05, **p < 0.01, ****p < 0.0001.

### Slide 7
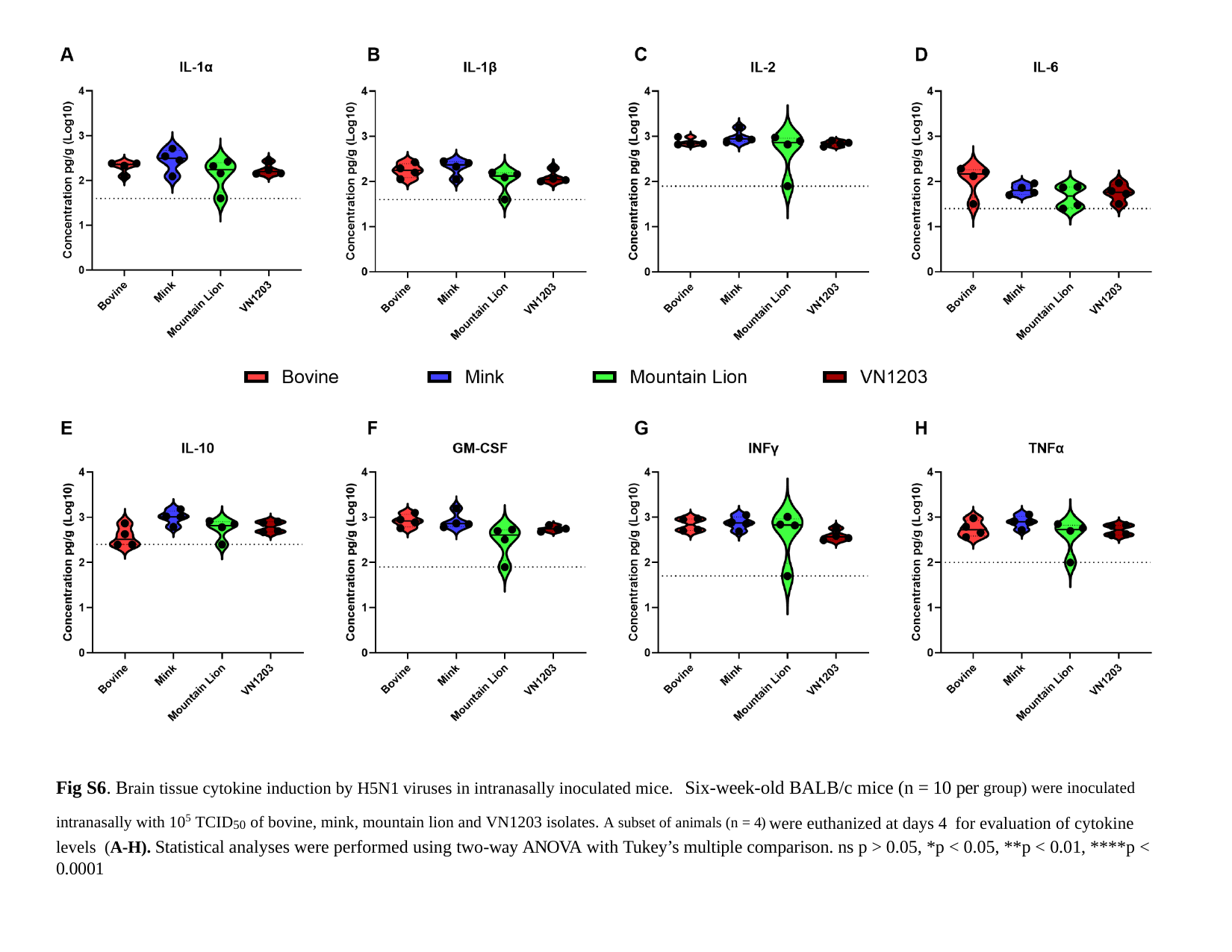

Fig S6. Brain tissue cytokine induction by H5N1 viruses in intranasally inoculated mice. Six-week-old BALB/c mice (n = 10 per ^group)^ ^were^ ^inoculated^ ^intranasally^ ^with^ ^105^ ^TCID^50 ^of^ ^bovine,^ ^mink,^ ^mountain^ ^lion^ ^and^ ^VN1203^ ^isolates.^ ^A^ ^subset^ ^of^ ^animals^ ^(n^ ^=^ ^4)^ were euthanized at days 4 for evaluation of cytokine levels (A-H). Statistical analyses were performed using two-way ANOVA with Tukey’s multiple comparison. ns p > 0.05, *p < 0.05, **p < 0.01, ****p < 0.0001

### Slide 8
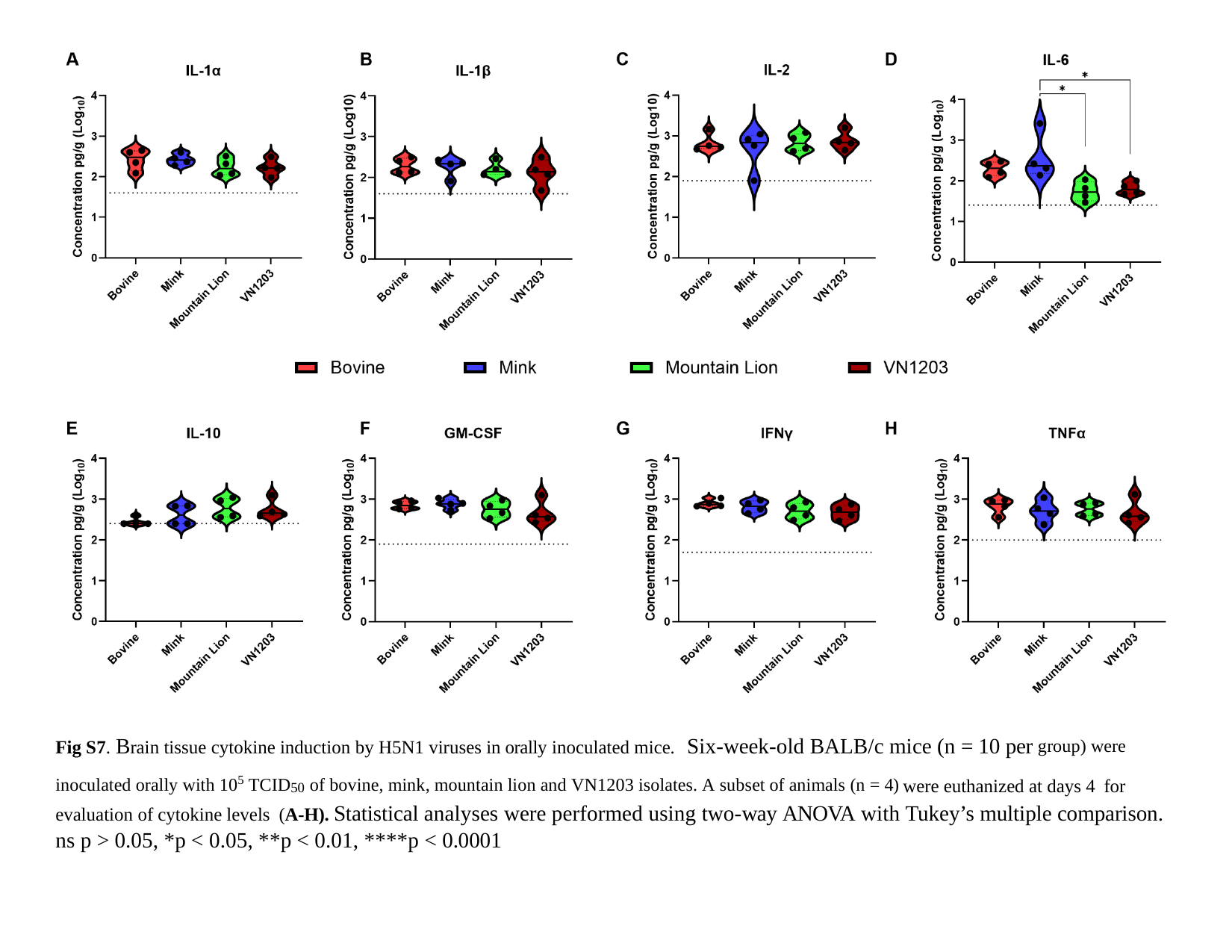

Fig S7. Brain tissue cytokine induction by H5N1 viruses in orally inoculated mice. Six-week-old BALB/c mice (n = 10 per ^group)^ ^were^ ^inoculated^ ^orally^ ^with^ ^105^ ^TCID^50 ^of^ ^bovine,^ ^mink,^ ^mountain^ ^lion^ ^and^ ^VN1203^ ^isolates.^ ^A^ ^subset^ ^of^ ^animals^ ^(n^ ^=^ ^4)^ were euthanized at days 4 for evaluation of cytokine levels (A-H). Statistical analyses were performed using two-way ANOVA with Tukey’s multiple comparison. ns p > 0.05, *p < 0.05, **p < 0.01, ****p < 0.0001

### Slide 9
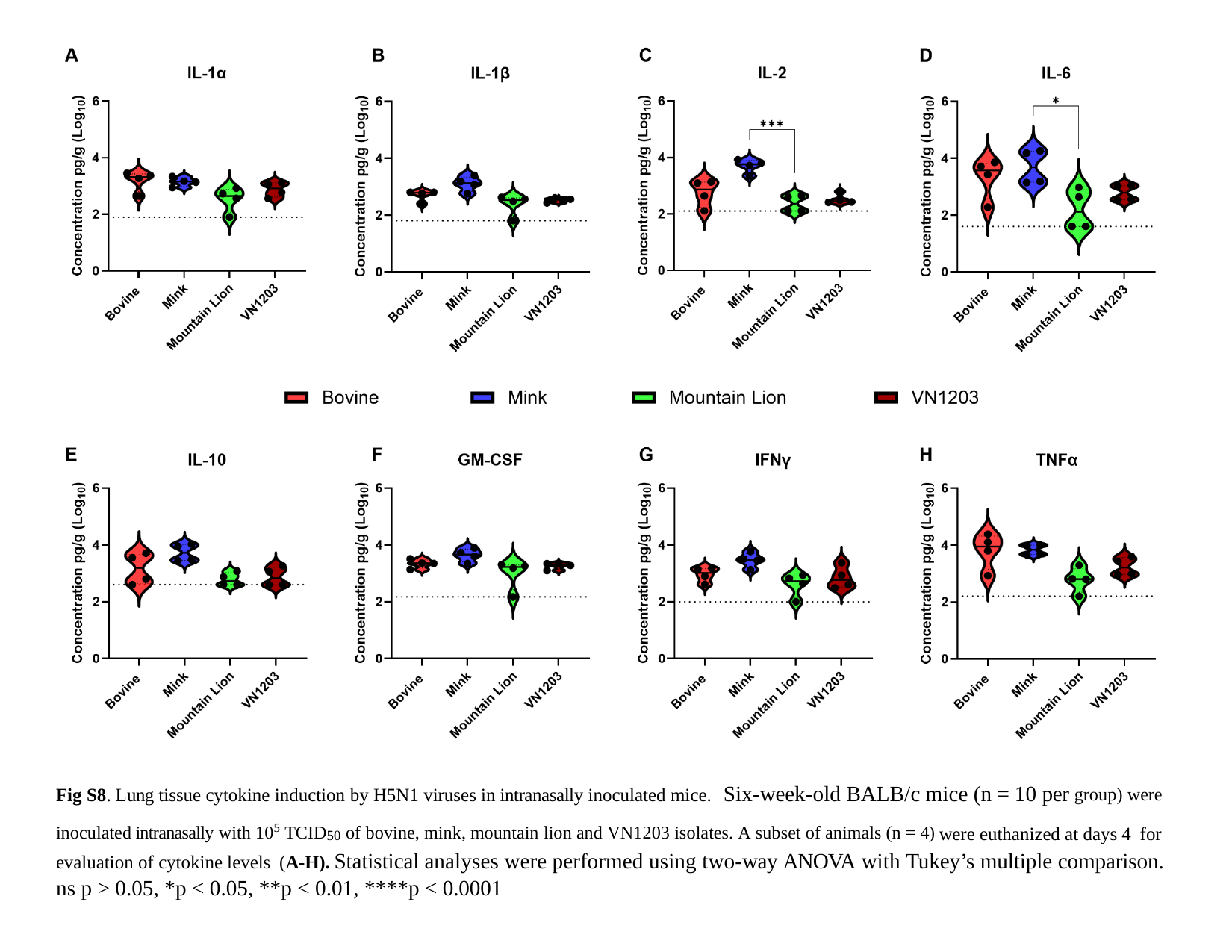

Fig S8. Lung tissue cytokine induction by H5N1 viruses in intranasally inoculated mice. Six-week-old BALB/c mice (n = 10 per ^group)^ ^were^ ^inoculated^ ^intranasally^ ^with^ ^105^ ^TCID^50 ^of^ ^bovine,^ ^mink,^ ^mountain^ ^lion^ ^and^ ^VN1203^ ^isolates.^ ^A^ ^subset^ ^of^ ^animals^ ^(n^ ^=^ ^4)^ were euthanized at days 4 for evaluation of cytokine levels (A-H). Statistical analyses were performed using two-way ANOVA with Tukey’s multiple comparison. ns p > 0.05, *p < 0.05, **p < 0.01, ****p < 0.0001

### Slide 10
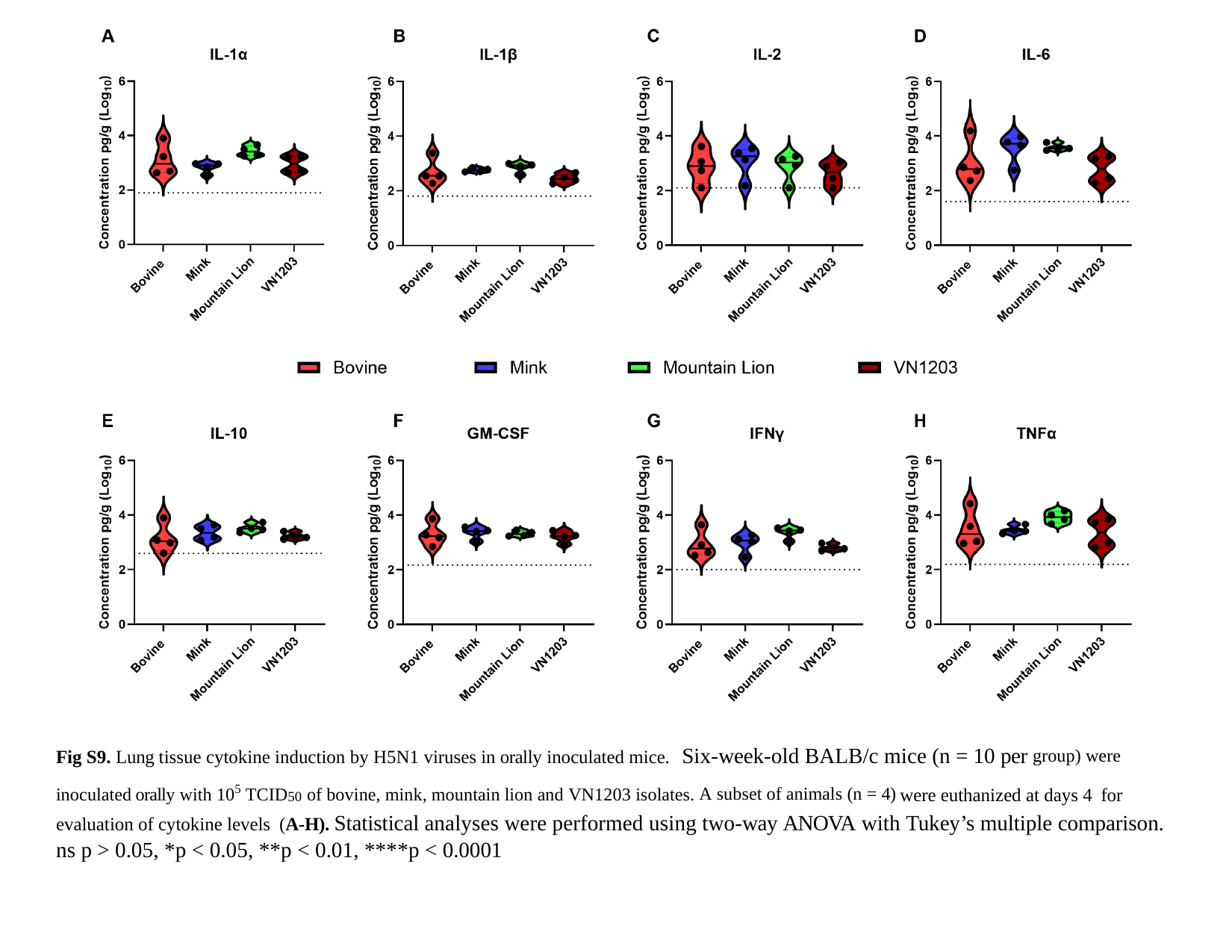

Fig S9. Lung tissue cytokine induction by H5N1 viruses in orally inoculated mice. Six-week-old BALB/c mice (n = 10 per ^group)^ ^were^ ^inoculated^ ^orally^ ^with^ ^105^ ^TCID^50 ^of^ ^bovine,^ ^mink,^ ^mountain^ ^lion^ ^and^ ^VN1203^ ^isolates.^ ^A^ ^subset^ ^of^ ^animals^ ^(n^ ^=^ ^4)^ were euthanized at days 4 for evaluation of cytokine levels (A-H). Statistical analyses were performed using two-way ANOVA with Tukey’s multiple comparison. ns p > 0.05, *p < 0.05, **p < 0.01, ****p < 0.0001
